# A thermodynamic model for water activity and redox potential in evolution and development

**DOI:** 10.1101/2021.01.29.428804

**Authors:** Jeffrey M. Dick

**Affiliations:** Key Laboratory of Metallogenic Prediction of Nonferrous Metals and Geological Environment Monitoring, Ministry of Education, School of Geosciences and Info-Physics, Central South University, Changsha 410083, China

**Keywords:** geobiochemistry, protein evolution, biofilm development, proteomics, water activity, oxidation state

## Abstract

Reactions involving water and oxygen are basic features of geological and biological processes. To understand how life interacts with its environment requires monitoring interactions with H_2_O and O_2_ at timescales relevant to not only organismal growth but also billions of years of geobiological evolution. Here, chemical transformations intrinsic to evolution and development were characterized by analyzing data from recent phylostratigraphic and proteomic studies. This two-stage analysis involves obtaining chemical metrics (carbon oxidation state and stoichiometric hydration state) from the elemental compositions of proteins followed by modeling the relative stabilities of target proteins against a proteomic background to infer thermodynamic parameters (oxygen fugacity, water activity, and virtual redox potential (Eh)). The main results of this study are a rise in carbon oxidation state of proteins spanning the time of the Great Oxidation Event, a rise in virtual redox potential that coincides with the likely emergence of aerobic metabolism, a rise in carbon oxidation state of proteins inferred from the transcriptome in late stages of *Bacillus subtilis* biofilm growth, and a drop in stoichiometric hydration state of the fruit fly developmental proteome at the same time as a drop in organismal water content. Stoichiometric hydration state also decreases for proteins with more recent gene ages and through stages of biofilm development, leading to predictions of higher hydration potentials at earlier time points. By building chemical representations of protein evolution and developmental proteomes, exciting new types of geobiochemical proxies can be developed.

## Introduction

Many examples of dynamic hydration levels can be found in biology. Water content is one of the most basic biochemical quantities that changes during development (Church and Robertson, 1966; Friis-Hansen, 1983), and water is progressively lost from prenatal to adult forms in mammals (Logan and Himwich, 1972; Calcagno et al., 1972; Friis-Hansen, 1983). Conversely, relatively high water content has been recognized for over a century as a biochemical characteristic of cancer tissue (Cramer, 1916; Downing et al., 1962; Saryan et al., 1974; Ross and Gordon, 1982), and some authors have highlighted parallel trends of higher water content in both cancer and embryonic tissue (Winzler, 1959; Olmstead, 1966; McIntyre, 2006). In origin-of-life research, geological environments with low water activity 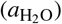 have been proposed to reduce or overcome the energetic barriers to polymerization of biomolecules in an aqueous environment (Pace, 1991). This concept has received renewed attention recently for serpentinizing systems, which could provide not only oxidation-reduction (redox) disequilibria to drive the abiotic synthesis of various organic molecules but also pore spaces with reduced water activity (Lamadrid et al., 2017).

The importance of both redox conditions and water activity for life’s origin in geological environments (do Nascimento Vieira et al., 2020), not to mention the fact that water is involved in far more biochemical reactions than any other metabolite (Frenkel-Pinter et al., 2021), raises the question whether biological processes in general are also shaped by these chemical parameters. With the growing availability of gene age estimates for evolution and measurements of protein abundance during organismal development, it it now possible to construct thermodynamic models for chemical transformations of the most abundant type of biomacromolecule – that is, proteins – that occur at timescales longer than cellular lifetimes.

Thermodynamic principles predict the most stable chemical species, but don’t say how the reactions that lead to them take place (Bard, 1986). Relative stability diagrams for minerals, also known as chemical activity or predominance diagrams, are immensely useful for interpreting geological phenomena, yet are agnostic about nucleation effects or whether the minerals grow through precipitation from solution or solid-state diffusion and replacement. This is because it is only the Gibbs energy of the system, and not any particular reaction mechanism, that determines the relative stabilities of chemical species.

In the same way that stability diagrams for mineral phases neither inform nor conflict with mechanistic models of nucleation or precipitation and replacement reactions, an analogous thermodynamic depiction of protein sequences does not conflict with elementary biochemical models of ATP conjugation and amino acid synthesis and polymerization. Such a model is likewise ambivalent about biological mechanisms at all levels from DNA replication and mutation to protein function, survival, and natural selection. By saying nothing about these mechanisms, the model independently predicts which proteins are more stable in a multidimensional space defined by temperature, pressure, and chemical potentials – the fundamental variables in thermodynamics.

The consideration of both water activity and oxygen fugacity 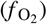 – a thermodynamic indicator of oxidation potential – is essential for thermodynamic models of melting and magmatic processes (Foley, 2011) as well as lower-temperature metasomatic processes including serpentinization (Evans et al., 2013). Although 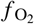may have no physical meaning as an indicator of partial pressure, it has a thermodynamic definition that makes it a useful indicator of the internal oxidation potential of systems without an actual gas phase (Frost, 1991). For instance, an oxygen fugacity given by log 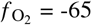 (where log denotes the common logarithm) is many orders of magnitude lower than an estimated upper bound of partial pressure of O_2_ in the prebiotic Archean atmosphere (*<* 10^−13^ bars; Kasting, 1993; Catling and Claire, 2005), and would correspond to one molecule of O_2_ in a volume approximately equal to that of the solar system (Anderson, 2005, p. 365). Despite its unphysical interpretation as a scale of partial pressure, 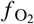can be converted to meaningful values of redox potential in the Eh scale. By way of example, for unit activity of water (that is, 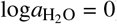), values of 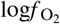between −40 and −83.1 correspond to Eh values between approximately −70 and −410 mV at pH = 7 and 25 °C (Garrels and Christ, 1965, p. 176).

Geochemistry is primarily concerned with reactions among minerals and dissolved aqueous species; geobiochemistry, as envisioned in this study, applies the same concepts to groups of proteins that constitute an evolutionary or developmental series. The thermodynamic model here is parameterized in terms of water activity and oxygen fugacity, which are then converted to the Eh scale of redox potential that is more common in biology. This method offers a path toward quantitative thermodynamic retrodiction of past redox conditions by comparing groups of proteins classified by the evolutionary ages of genes (phylostrata). Similarly, a chemical representation of developmental patterns of protein expression provides novel insight into the role of water at the nexus between protein expression and cell physiology.

## Materials and Methods

### Chemical metrics

Two chemical metrics derived from the elemental composition of proteins are used in this study – carbon oxidation state and stoichiometric hydration state. This chemical representation is the precursor to a thermodynamic analysis that provides a theoretical assessment of intensive chemical potentials, including water activity.

Carbon oxidation state (*Z*_C_) represents the average charge on carbon atoms in a molecule, given nominal charges of the other atoms (H^+1^, N^-3^, O^-2^, S^-2^). The carbon oxidation state can be computed directly from the elemental abundances of molecules in which the heteroatoms (N, O, S) are bonded only to H and/or C but not to each other (Dick, 2014; Dick et al., 2020). This condition holds for the 20 standard amino acids and for primary sequences of proteins but not for disulfide bonds and some types of post-translational modifications, none of which are considered here.

Stoichiometric hydration state 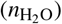 is the coefficient on H_2_O in the mass-balanced reaction representing the theoretical formation of a protein from a set of thermodynamic components, also known as basis species. The basis species glutamine, glutamic acid, cysteine, H_2_O, and O_2_ (denoted “QEC”) were chosen for this analysis (Dick et al., 2020; Dick, 2021b). Any choice of basis species is permissible, but only one set was used for all the calculations here. The particular choice of basis species was made to (1) strengthen the covariation between two measures of oxidation state: *Z*_C_ (which does not depend on the choice of basis species) and the number of O_2_ in the formation reactions from basis species, and (2) reduce the covariation between *Z*_C_ and 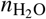(see Dick et al., 2020 for details). Therefore, *Z*_C_ and 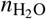can be regarded as largely uncoupled variables that enable changes of oxidation and hydration state to be identified as nearly perpendicular trends on 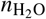*–Z*_C_ scatterplots.

In order to represent chemical differences associated with amino acid composition, values of 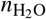for individual proteins were normalized by protein length. To illustrate the necessity for normalization, a 200-aa protein has more O and H atoms than a 100-aa protein, so the number of H_2_O molecules in the theoretical formation reaction is greater for the larger protein. Normalization makes it so comparisons of 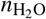are not dominated by protein length, but instead represent differences in amino acid composition. In contrast, because *Z*_C_ is defined as a per-carbon quantity, it does not scale with protein length, and normalizing it by protein length would be inappropriate.

Chemical metrics (*Z*_C_ and 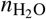) for individual proteins were used to calculate mean values and bootstrap confidence intervals for groups of proteins. The calculations were performed either without weighting (for phylostratigraphic datasets or proteomic datasets that list up- and down-regulated proteins) or with weighting by abundance (for proteomic or transcriptomic datasets that give expression levels). Means of chemical metrics were not weighted by protein length, so large and small proteins contribute equally to the mean value. To calculate thermodynamic parameters (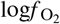and 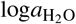), each group of proteins was modeled by a single target protein having the mean amino acid composition of the group. In contrast to the mean values for chemical metrics, the mean amino acid compositions for target proteins include weighting by both abundance (if available) and implicitly by protein length, as longer proteins have more amino acid residues.

### Data sources

Phylostrata were obtained from the supporting information of Trigos et al. (2017) and the “main HUMAN.csv” file of Liebeskind et al. (2016a,b). The Ensembl gene identifiers in the Trigos et al. dataset were converted to UniProt accession numbers (The UniProt Consortium, 2019) using the UniProt mapping tool (Huang et al., 2011); 169 of 17,318 genes could not be mapped to UniProt and were removed from the subsequent analysis.

Transcriptomic and proteomic data for growing *Bacillus subtilis* biofilms were taken from Supplementary file S10 of Futo et al. (2021), specifically the tables named “Input values for calculating TAI” and “Input values for calculating PAI”. The phylostrata given by Futo et al. (2021) were not included in this analysis. Data for the developmental proteome of *Drosophila melanogaster* were extracted from Supplemental Table S1 of Casas-Vila et al. (2017). The values in the columns for imputed log_2_ LFQ intensity were exponentiated, then mean values were computed for each time point (4 replicates). For both the *B. subtilis* and *Drosophila* datasets, protein IDs were mapped using the UniProt mapping tool (Huang et al., 2011). The canonical protein sequences were downloaded from UniProt, and the read.fasta() function in the CHNOSZ package (Dick, 2019) was used to compute the amino acid compositions of the proteins.

Differentially expressed proteins between embryos and adult flies were obtained from Supplementary Table S2 of Fabre et al. (2019). After rounding the values in the “Log2 ratio Adult/Embryo” column to 2 decimal places and those in the “-log10 BH corrected p-value Adult vs Embryo” column to whole numbers, the table was filtered to include proteins with log_2_ expression change *≥* 1 or *≤* −1 and log_10_ *p*-value *≤* 2. This yielded 407 and 369 proteins with higher expression in adults and embryos, respectively; these are very close to the numbers given by Fabre et al. (2019) (407 and 371 proteins enriched in adults and embryos).

### Computer code

All figures were created in R version 4.1.0 (R Core Team, 2021). The R package CHNOSZ version 1.4.1 (Dick, 2019) was used for thermodynamic calculations, and canprot version 1.1.0 (Dick, 2021b) was used for calculating chemical metrics from amino acid compositions of proteins. The R package boot version 1.3-28 (Canty and Ripley, 2021; Davison and Hinkley, 1997) was used for calculating bootstrap confidence intervals (95% confidence levels of the percentile type for 999 bootstrap replicates).

## Results

### Scope and limitations of phylostratigraphic datasets

Phylostrata represent conservation levels of orthologous genes in a phylogenetic tree (Domazet-Lošo et al., 2017; Van Oss and Carvunis, 2019). Although the complex sequence of evolutionary events for any gene precludes a single, well-defined age, an operational definition used in phylostratigraphy considers the presence or absence of a gene family in different species (i.e. Dollo parsimony), leading to the inference that the gene family originated in the most recent common ancestor of all species that have that gene family (Capra et al., 2013). Accordingly, applications of phylostratigraphic analysis often refer to the evolutionary age of particular genes (Trigos et al., 2017; Zhou et al., 2018; Futo et al., 2021). However, true orthologs that cannot be detected (false negatives) become more common with greater evolutionary divergence (Natsidis et al., 2021), which adds to the uncertainty of phylostratigraphic age estimates.

Homology detection based on protein domains can reduce false negatives compared to that based on full gene sequences. Sensitivity can also be increased by using hidden Markov models for multiple sequences (instead of pairwise sequence alignment), and statistical power is enhanced by analyzing the full genomes of hundreds of species rather than particular focal species. By using these techniques, James et al. (2021) found that the support for phylostratigraphic trends in protein hydrophobicity and intrinsic structural disorder was strengthened compared to previous studies. Accordingly, although false negatives may have been more prevalent in earlier phylostratigraphic studies, they are probably random errors that do not create spurious trends, but simply reduce the power to detect trends. It was also noted that ancient domains are strongly influenced by the amino acid availability at the time of *de novo* gene birth, even after billions of years of evolution (James et al., 2021). The preservation of ancient patterns of amino acid composition in contemporary sequences supports the approach used in the present study of comparing the chemical metrics for proteins in different age groups to geological events.

A further limitation is that phylostratigraphy represents ages of genes in a particular species (Natsidis et al., 2021). Therefore, phylostrata divide up a single modern proteome into age groups and do not yield complete ancestral proteomes. For human genes, the number of genes assigned to particular phylostrata varies widely (Fig. 1); in the dataset of Trigos et al. (2017) there are as many as 4665 for PS 2 (Eukaryota) and as few as 25 for PS 16 (*Homo sapiens*) (these counts exclude 10 genes in each of those phylostrata that could not be mapped to UniProt IDs). Therefore, the chemical analysis performed here only characterizes subsets of proteins associated with particular phylogenetic branches, and not entire proteomes.

**Fig. 1.**
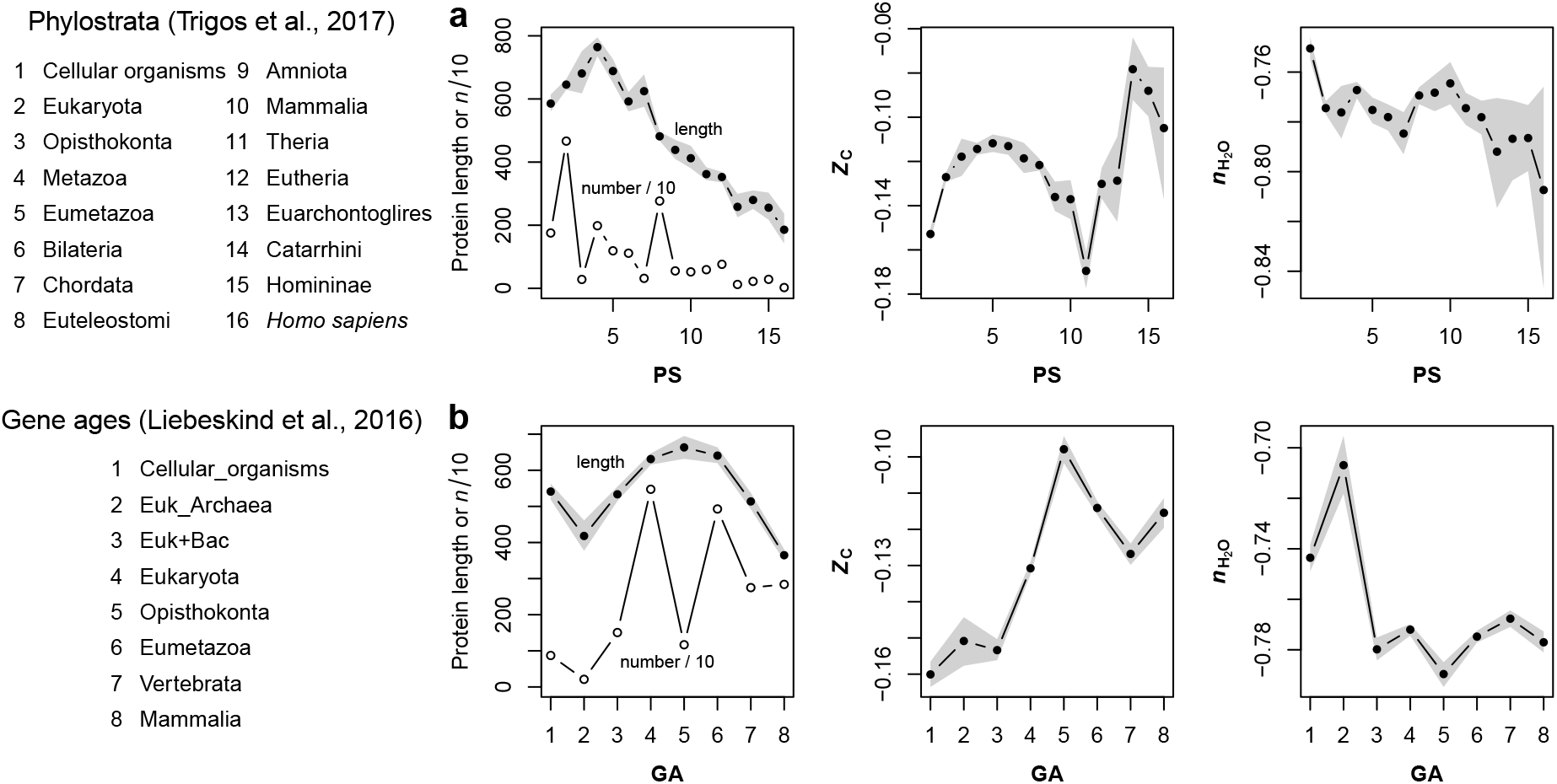
Protein length and chemical metrics for phylostratigraphic age groups. **a** Mean values of protein length, *Z*_C_, and 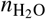of proteins for all protein-coding genes in each phylostratum (PS) given by Trigos et al. (2017). The points represent the mean values for individual phylostrata, and the shaded areas represent bootstrap confidence intervals. In the first plot, the (number of proteins in each phylostratum) / 10 is plotted on the same scale as protein length. **b** The same analysis for proteins with gene ages (GA) given by Liebeskind et al. (2016b). The last age group in this dataset is Mammalia, which corresponds to PS 10 in the Trigos et al. dataset

Another limitation of this analysis is that actual expression levels vary dramatically for different proteins, but phylostratigraphic datasets do not provide any abundance information. In the second part of this study, protein expression levels are used to construct abundance-weighted means of chemical metrics and amino acid composition for developmental proteomes. Such an abundance-weighted mean is not possible for the chemical and thermodynamic analysis of phylostratigraphic data.

### Chemical analysis of proteins in phylostratigraphic age groups

The mean lengths of proteins coded by genes in each of 16 phylostrata (PS) for human protein-coding genes given by Trigos et al. (2017) are plotted in Fig. 1a. There is an initial rise in protein length leading up to Eukaryota, which is consistent with the previously reported greater median protein length in eukaryotes than prokaryotes (Brocchieri and Karlin, 2005). Although it is debated whether the large decline of protein length for younger genes (i.e. those with lower conservation levels) is real (Lipman et al., 2002; Domazet-Lošo et al., 2017) or to some extent an artifact of BLAST-based homology detection (Moyers and Zhang, 2017; Van Oss and Carvunis, 2019), controlling for length either strengthens or has no effect on inferred trends of intrinsic structural disorder (ISD) (Wilson et al., 2017; Heames et al., 2020). ISD predicted from amino acid sequences of proteins is strongly associated with hydrophobicity (James et al., 2021); similarly, the chemical metrics used here are derived from the amino acid compositions of proteins, and *Z*_C_ exhibits a negative correlation with amino acid hydrophobicity (Dick, 2014), so trends in these metrics are likely to be robust against length-dependent artifacts.

Figure 1a reveals distinct evolutionary patterns of oxidation state and hydration state of proteins. *Z*_C_ forms a strikingly smooth hump between PS 1 and 11 then increases rapidly to the maximum at PS 14, followed by a smaller decline to PS 16, which corresponds to *Homo sapiens*. In contrast, 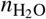shows an overall decreasing trend in later phylostrata, although there are positive jumps between PS 3 and 4 and PS 7 and 8.

Figure 1b shows the same analysis applied to proteins grouped into eight gene ages (GA) reported by Liebeskind et al. (2016b) based on consensus tables of gene-age estimates from different orthology inference algorithms. The gene ages of Liebeskind et al. have three steps between cellular organisms and Eukaryota, providing a greater resolution in earlier evolution compared to the phylostrata of Trigos et al., and stop at Mammalia, which corresponds to PS 10 of Trigos et al.. Keeping in mind the different resolutions and scales of the Trigos et al. phylostrata and Liebeskind et al. gene ages, the two datasets show similar maxima for *Z*_C_ and protein length between the the origins of Eumetazoa and Opisthokonta, and an overall decrease of 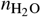in younger age groups (Fig. 1b).

### Thermodynamic model for maximum activities of proteins

To take the step from chemical composition to chemical potential, a thermodynamic model is described here for predicting the oxidation and hydration potentials that stabilize particular proteins compared to others. This analysis of relative stability does not refer to the forces that determine protein conformations (i.e. 3-dimensional structures), but to the Gibbs energies of the theoretical formation reactions of proteins from basis species. It follows that the relative stabilities of proteins are influenced by the chemical activities of basis species whose stoichiometric coefficients depend on the elemental abundances and therefore amino acid sequences of different proteins.

To give a worked-out example, the balanced formation reaction of a well-known protein, chicken egg-white lysozyme (UniProt: P00698, LYSC CHICK), can be written as:

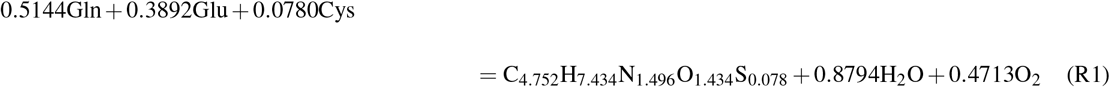

Here, the chemical formula of the whole protein (C_613_H_959_N_193_O_185_S_10_) is divided by the protein length (129) to give the per-residue formula that is a product of the reaction. The only other species in the reaction are the basis species glutamine (C_5_H_10_N_2_O_3_), glutamic acid (C_5_H_9_NO_4_), cysteine (C_3_H_7_NO_2_S), H_2_O, and O_2_. The coefficient on H_2_O in this reaction is the opposite of the stoichiometric hydration state; that is, 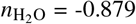. The carbon oxidation state of the protein, *Z*_C_ = 0.016, can be computed from either its chemical formula or amino acid composition (see Dick et al., 2020).

The central thermodynamic quantity in this model is chemical affinity (*A*), which is the opposite of the non-standard Gibbs energy change of the reaction (Denbigh, 1981, p. 143) and can be computed from (e.g. Solel et al., 2019)

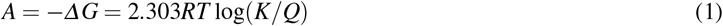

where *K* is the equilibrium constant and *Q* is the activity product for the formation reaction for a particular protein. Factorization using the natural logarithm of 10 (*≈*2.303) means that common logarithms of variables are used throughout. Because it includes the chemical activities of all the species in the reaction, *Q* is affected by both 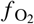and 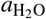, which represent oxidation and hydration potential. On the other hand, *K* is a function of the standard Gibbs energy of the reaction and therefore of temperature and pressure.

The standard Gibbs energies (Δ*G*_f_°) of proteins were calculated using amino acid group additivity as described by Dick et al. (2006) together with later updates for methionine and glycine groups and the protein backbone (LaRowe and Dick, 2012; Kitadai, 2014). For the purpose of quantifying the relative stabilities of proteins in terms of hydration and oxidation potential, the calculations were limited to proteins treated as neutral species using values of Δ*G*_f_° calculated for 25 °C and 1 bar. For each protein, Δ*G*_f_° was also divided by the protein length to give the per-residue value, which was combined with the standard Gibbs energies of the other species in the reaction to calculate the standard Gibbs energy of reaction (*G*_r_°), and from that, log*K*. The standard Gibbs energies for H_2_O, O_2_, and amino acids were taken from Haar et al. (1984), Wagman et al. (1982), and Dick et al. (2006). By using the subcrt() function in the CHNOSZ package (Dick, 2019), log*K* for Reaction (R1) at 25 °C and 1 bar can be computed to be −39.84.

Calculation of the activity product (*Q*) requires values for activities of all the species in the reaction; because oxygen is a gas, the fugacity of O_2_ is used instead of activity. 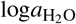 and 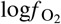were chosen as exploratory variables, so their values correspond to the axes of a 2-dimensional grid (256×256 resolution) used to plot the relative stabilities of the proteins. Although the chemical activities of all the basis species could be considered as variables in the thermodynamic model, it is not feasible to produce a 5-dimensional visualization of the system. Because 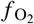and 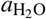 were identified as the major variables of interest, the activities of the others were set to constants that correspond to biochemically reasonable concentrations. Specifically, mean concentrations of amino acids in human plasma (Tcherkas and Denisenko, 2001) were used; expressed as logarithms of concentrations in mol/l, these are −3.2 for glutamine, −4.5 for glutamic acid, and −3.6 for cysteine. Finally, the activity of the per-residue formula for each protein was set to unity. The activities were combined to give values of *Q* that were then used to calculate chemical affinity from Eq. (1). Continuing the example for LYSC CHICK, log*Q* for Reaction (R1) at nominal values of 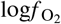(−70) and 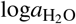 (0) is −29.33, which yields log(*K*/*Q*) = −10.52.

The relative stabilities of proteins represented by their per-residue formulas were computed by using the Boltzmann distribution written as

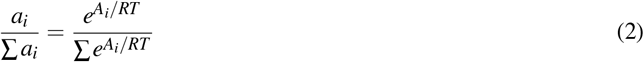

where *a* is activity, *A* is affinity calculated given unit activity of the per-residue formula for each protein, and *i* designates a single protein in a system of any number of proteins. Activity coefficients were taken to be unity, and the total activity (∑ *a*_*i*_) was also fixed at unity. Consequently, *a*_*i*_ for each protein represents its fractional abundance at equilibrium with all other proteins in the system.

According to the equilibrium model, each protein in a system of candidate proteins has a finite value of activity that depends on 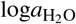 and 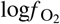, which are the only free variables. The predominant protein for any combination of these variables (“condition”) is the one with the highest predicted activity. In thermodynamic models for inorganic systems, analogous calculations are used to make predominance diagrams for aqueous species (e.g. Kinniburgh and Cooper, 2004), but for the analysis of biological data it is more informative to find the sets of conditions that respectively maximize the activity of each of the proteins in a system and not only the predominant one. This model, which is referred to as “MaximAct” in the present study, is described next.

### Maximizing stabilities of target proteins on a proteomic background

In order to thermodynamically characterize the differences in protein composition between phylostrata, model proteins for each phylostratum (referred to here as target proteins) were generated by computing the mean amino acid composition of all proteins in each phylostratum. Equilibrium calculations for the 16 target proteins for the Trigos et al. phylostrata are displayed on the 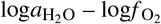 diagram in Fig. 2a. Predominance fields for just a few proteins are visible on the diagram. These fields represent the proteins that have the highest activities; those with lower activity by definition do not predominate, but they still have calculable values of chemical activity. The conditions for maximum activity for each of the 16 target proteins (indicated by the points) are found at at extreme values of 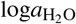 and 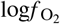. Finite conditions for maximum activity of some of the proteins could be found by extending the range of the diagram, but the activities of the predominant proteins maximize at infinite values of 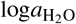 and/or 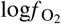. Therefore, it is not possible to use this simple model to find particular values of water activity and oxygen fugacity that characterize each phylostratum.

**Fig. 2.**
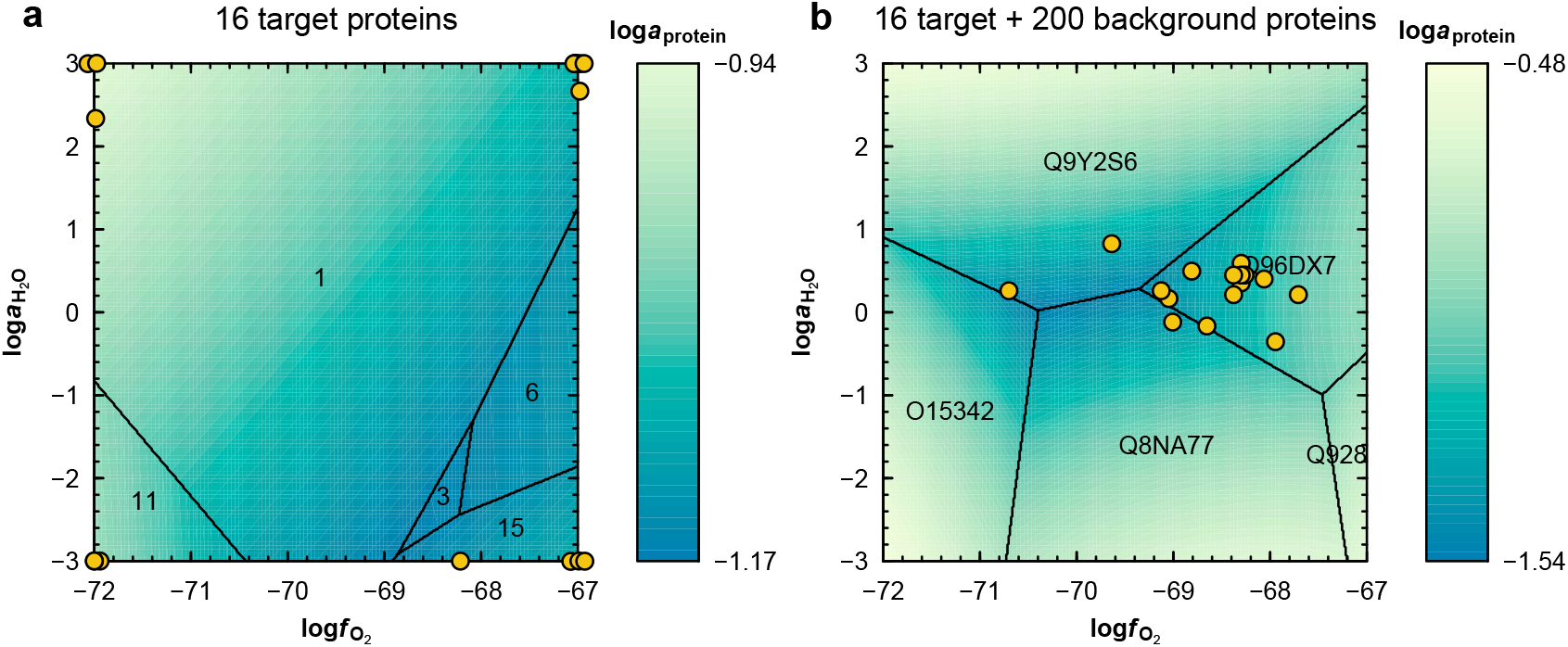
Strategy for deriving values of water activity (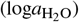) and oxygen fugacity (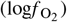) that maximize the predicted equilibrium activity of target proteins defined by the mean amino acid compositions for phylostrata. **a** A system consisting of 16 target proteins derived from the phylostrata of Trigos et al. (2017). The stability fields represent the computed predominant proteins at equilibrium and the lines represent equal activities for the predominant proteins. Only five target proteins predominate in the plotted range of 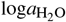 and 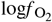; all the others have lower activity. The yellow circles indicate the conditions for the maximum activity of each of the target proteins; a small amount of jitter is added to aid visualization. **b** A system that includes the same 16 target proteins and 200 randomly sampled proteins from the human proteome (background proteins). In this case, the predicted predominant proteins all come from the background population, and are labeled with their UniProt IDs. The target proteins have lower activities that are maximized at the conditions indicated by the yellow circles

A key innovation in this study is the addition of more proteins to the thermodynamic system to represent a background population, or biological context, that allows the transformations between target proteins to be resolved in finer detail. An appropriate set of background proteins would have a wider compositional range than the mean amino acid compositions used to make the target proteins. Because the background proteins are now competing for formation with the target proteins, the conditions for maximum activity of the latter will be constrained to smaller and more meaningful ranges of 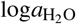 and 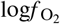. The background proteins should also represent the actual biological range of amino acid composition, so a plausible choice is to use all proteins coded by the human genome. To reduce model bias that could be caused by the inclusion of background proteins with unusual amino acid compositions, a representative human proteome background was constructed by taking the intersection of UniProt IDs for proteins coded by genes in the Trigos et al. and Liebeskind et al. datasets, yielding 16,723 unique sequences.

The calculated predominance diagram for a system of 16 target proteins together with 200 randomly sampled background proteins reveals that a relatively small number of background proteins predominate at equilibrium (Fig. 2b). None of the 16 target proteins predominates, but instead their activities are maximized at particular values of 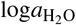 and 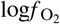. Figure 2b is a result that corresponds to only one random subsample of the background human proteome. There are many possible versions of Fig. 2b, which are not shown here. Instead, the 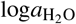 and 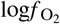values obtained by using many random subsamples of the human proteome with a larger number of sampled proteins (2000) are shown in Fig. 3a and b. This number was chosen to be able to run the calculation within computer memory limits, and the random subsampling was repeated 100 times to obtain mean values for 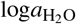 and 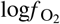, which approximate the values that would be obtained if a single calculation could be performed with all 16,723 proteins in the background proteome.

**Fig. 3.**
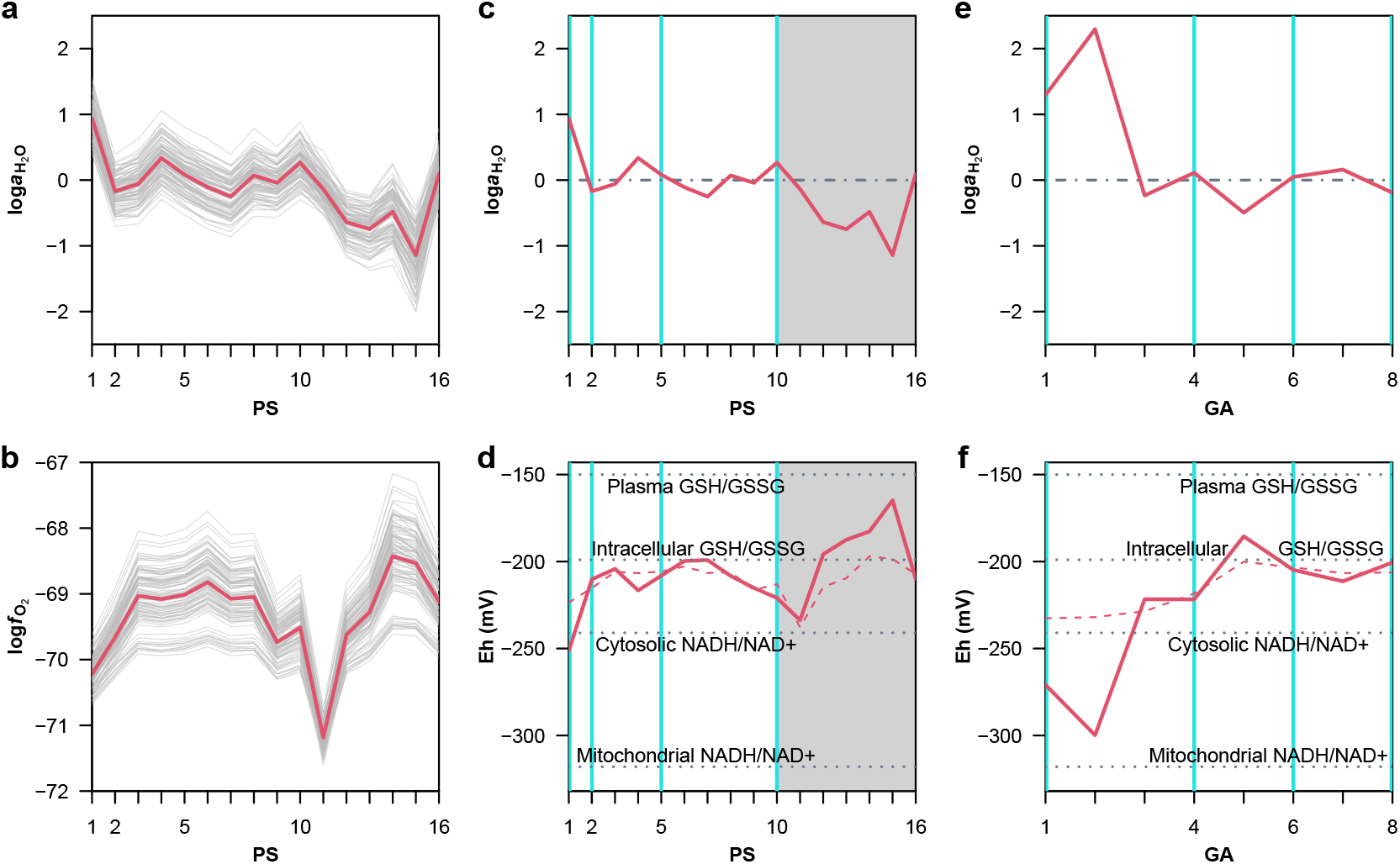
Analysis of optimal water activity and oxygen fugacity for phylostrata, and calculation of virtual redox potential (Eh). **a** and **b** Values of 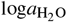 and 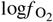that maximize the activity of target proteins for the Trigos et al. phylostrata in equilibrium with each other and 2000 randomly sampled background proteins. Each thin gray line represents a calculation for one random sample, and thick red lines show the means for 100 calculations. **c** Comparison of mean values of 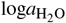 with pure water (dot-dashed horizontal line at [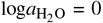). Vertical lines at cellular organisms, Eukaryota, Eumetazoa, and Mammalia indicate coinciding gene ages in the Liebeskind et al. dataset. The shaded gray area represents phylostrata that postdate the emergence of Mammalia, which are not available in the Liebeskind et al. dataset. **d** Virtual Eh calculated from the mean values of 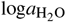 and 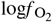using Eqs. (3)–(4) (thick red line) and from the same values of 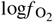with 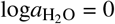 (dashed red line). Dotted horizontal lines represent redox potentials for different cellular compartments and reactions (van ‘t Erve et al., 2013; Jones and Sies, 2015). **e** and **f** Analogous calculations for 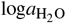 and Eh using the Liebeskind et al. gene ages; vertical lines represent cellular organisms, Eukaryota, Eumetazoa, and Mammalia

The variability among calculations displayed in Fig. 3a and b is due to the limited subsample size of background proteins, not to intrinsic biological variability. The use of a single mean amino acid composition for each target protein precludes the calculation of confidence intervals for thermodynamic parameters (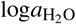 and 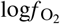), but relative uncertainties within datasets may be assessed by comparison with the confidence intervals for the corresponding chemical metrics.

It is not surprising to find that the trends of 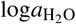 and 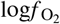depicted in Fig. 3a and b are similar to those of the corresponding chemical metrics, 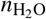and *Z*_C_, in Fig. 1a. For instance, PS 1 (cellular organisms) has both the highest 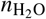and 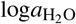. It is noteworthy that the water activity for PS 2–10 predicted by the MaximAct model fluctuates around unity, which is indicated by the horizontal dot-dashed line at 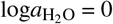 in Fig. 3c. Between PS 10 (Mammalia) and PS 15 (Homininae), both 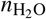and 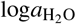 decrease considerably. However, although PS 16 (*Homo sapiens*) has the lowest 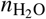of any phylostratum (Fig. 1a), the thermodynamic model predicts a higher 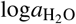 – closer to 0 – for PS 16 compared to PS 15. Looking at the oxidation trends, the broad hump in *Z*_C_ between PS 1 and 11 is reflected in 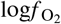values but with a more rugged profile, and PS 11 (Theria) has both the lowest *Z*_C_ and lowest 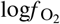(Fig. 3b).

### Virtual redox potential

Oxygen fugacity is a thermodynamic quantity that implies nothing about the actual mechanism of oxidation or reduction (Anderson and Crerar, 1993) and in practice is used to calculate other parameters that are easier to measure (Anderson, 2005, p. 245). In this study, the theoretical values of not only 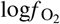but also 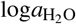 are used to calculate a virtual redox potential (Eh), which can be compared with independent measurements of Eh in biology. The qualifier “virtual” signifies that the values of Eh are not obtained from standard laboratory techniques, but rather from an independent thermodynamic model for the chemical differences of proteins over evolutionary and developmental time scales.

Virtual redox potential was calculated by considering the half-cell reaction for H_2_O:

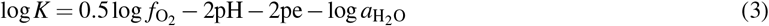

For equilibrium of Reaction (R2), we can write

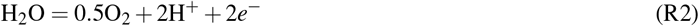

where pH = *−* log *a*_H_+ and pe = *−* log *a*_*e*_*−*. Eq. (3) was combined with pH = 7 and values of 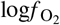and 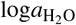 from the MaximAct model to calculate pe, which was then converted to Eh using

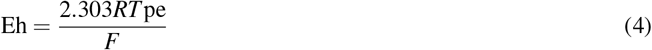

where *R, T*, and *F* are the gas constant, temperature in Kelvin, and Faraday constant.

Fig. 3d shows virtual Eh for the target proteins calculated using Eqs. (3)–(4) at 25 °C, 1 bar, and pH = 7. Virtual Eh is elevated by either increasing 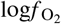or decreasing 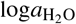. Compared to that of 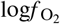, the whole Eh profile is tilted up, which reflects the decline of 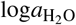 for proteins in more recent phylostrata. Several measurements for selected redox pairs in cells and plasma are shown for comparison (van ‘t Erve et al., 2013; Jones and Sies, 2015). The PS 1–11 hump begins and ends close to the Eh of the cytosolic NADH/NAD+ redox pair and maximizes near the Eh for intracellular GSH/GSSG measured in erythrocytes. Between PS 11 and 15 there is a rapid rise toward Eh values characteristic of GSH/GSSG in plasma, followed by a return in PS 16 to the redox potential for intracellular GSH/GSSG.

Figures 3e and f show the results of the MaximAct analysis applied to target proteins for the consensus gene ages of Liebeskind et al. (2016b). The first four gene ages correspond to the emergence of cellular organisms (GA 1), the last common ancestor of Eukaryota and Archaea (GA 2; Euk Archaea), genes that are present only in Eukaryotes and Bacteria, representing the horizontal transfer of genes to Eukaryotes (GA 3; Euk+Bac), and Eukaryota (GA 4). There is an increase of both 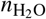(Fig. 1b) and 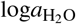 between GA 1 and 2, and these values are higher than those for all later gene ages. The target proteins for later gene ages are relatively stable near 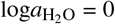 and are characterized by virtual Eh values that mostly range between cytosolic NADH/NAD_+_ and GSH/GSSG (Fig. 3f).

### Sensitivity of thermodynamic parameters to choice of proteomic background

It was argued above that the background proteome should reflect the complete range of biological amino acid composition. However, it might also be prudent to select the background proteins from the genome of the particular organism that is being studied. To examine these effects, Fig. 4a shows the chemical metrics for the proteomes of organisms considered in this study (human, *B. subtilis*, and *D. melanogaster*). Overlaid on this figure are the chemical metrics for target proteins for phylostrata and biofilm and fly development. The developmental proteomic datasets are analyzed separately below. Figure 4a is important because it shows that the target proteins for all datasets occupy a much smaller region of compositional space than the background proteins. This happens because the background proteins are individual protein sequences from a particular genome, while the target proteins have the mean amino acid compositions of groups of proteins identified in distinct phylostratigraphic and proteomic datasets. Therefore, the background proteins have a buffering effect that keeps the target proteins away from the edges of stability diagrams like that in Fig. 2b. Introducing a set of background proteins in relative stability calculations is a novel way of representing the internal milieu of cells, which is tightly regulated and has a restricted range of chemical properties compared to the exterior (Smith and Morowitz, 2016).

**Fig. 4.**
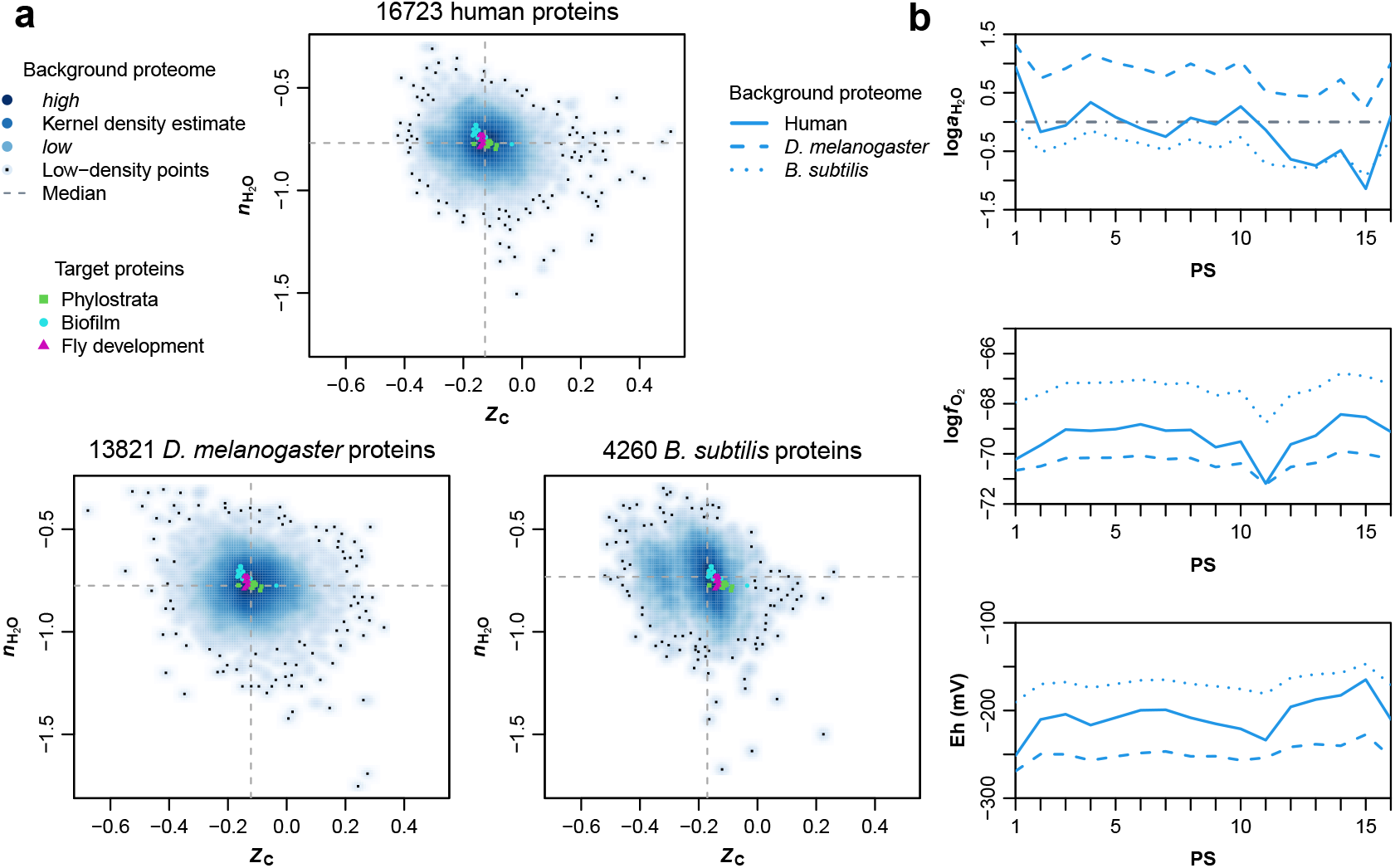
Chemical metrics and thermodynamic parameters calculated using different background proteomes. **a** Chemical metrics for background proteomes. The human background proteome was obtained as described in the text. *Drosophila melanogaster* and *Bacillus subtilis* proteomes were obtained from UniProt reference proteomes (The UniProt Consortium, 2019) for IDs UP000000803 and UP000001570 downloaded on 2021-07-12. Chemical metrics were calculated for individual protein sequences and plotted using the smoothScatter() function in R to generate a smoothed kernel density estimate of the data represented by color intensity. Small black symbols indicate the first 100 points in the low-density areas. Target proteins described in the text are indicated by differently colored symbols. The purpose of this plot is to show that the compositional range of all the target proteins for different datasets analyzed in this study is much smaller than that of the background proteomes; the differences between sets of target proteins are not analyzed here. **b** Thermodynamic parameters for target proteins for 16 phylostratigraphic age groups (based on Trigos et al., 2017) calculated using the MaximAct model with different background proteomes. For all other figures in this paper, the background proteome was obtained from the same organism as the source of target proteins.

Figure 4b shows how changing the background proteome for a single set of target proteins affects the thermodynamic parameters calculated with the MaximAct model. For instance, note that the *B. subtilis* proteome has a somewhat lower median carbon oxidation state and higher median stoichiometric hydration state compared to the human proteome (Fig. 4a). Because of the greater number of high-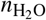proteins in the genome of *B. subtilis*, it provides background proteins that are relatively stable in the high-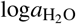 region of chemical potential space, which pushes the maximum activities of the target proteins to lower 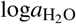 values (Fig. 4b). Likewise, the relatively low oxidation state of the *B. subtilis* proteome pushes the maximum activities of the target proteins to higher 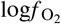values.

Changing the background proteome affects the precise values of thermodynamic parameters, but does not greatly affect the qualitative trends within a given dataset (Fig. 4b). For all other calculations in this study, the organism used for the background proteome was selected to be the same as that of target proteins. As a matter of future work, to assess the relative stabilities of target proteins among different organisms, a generic background proteome should be devised that is representative of the total range of biological amino acid composition and that would also be applicable to other domains of life.

### Chemical and thermodynamic analysis of biofilm development

In a transcriptomic and proteomic study of the development of *Bacillus subtilis* biofilms, Futo et al. (2021) described three periods of biofilm growth: early (6 hours to 1 day), mid (3 to 7 days), and late (1 to 2 months). Time points of 2 days and 14 days are regarded as transitional stages between these periods. In the early period of development, there is a decline in the mean protein length (Fig. 5a). This was computed by combining the lengths of canonical protein sequences from the UniProt database with normalized gene or protein expression values reported by Futo et al. (2021); no phylostrata assignments were used for this or any of the following calculations. The late period of biofilm development, for which only transcriptomic data are available, shows another drop in mean length of the corresponding proteins.

**Fig. 5.**
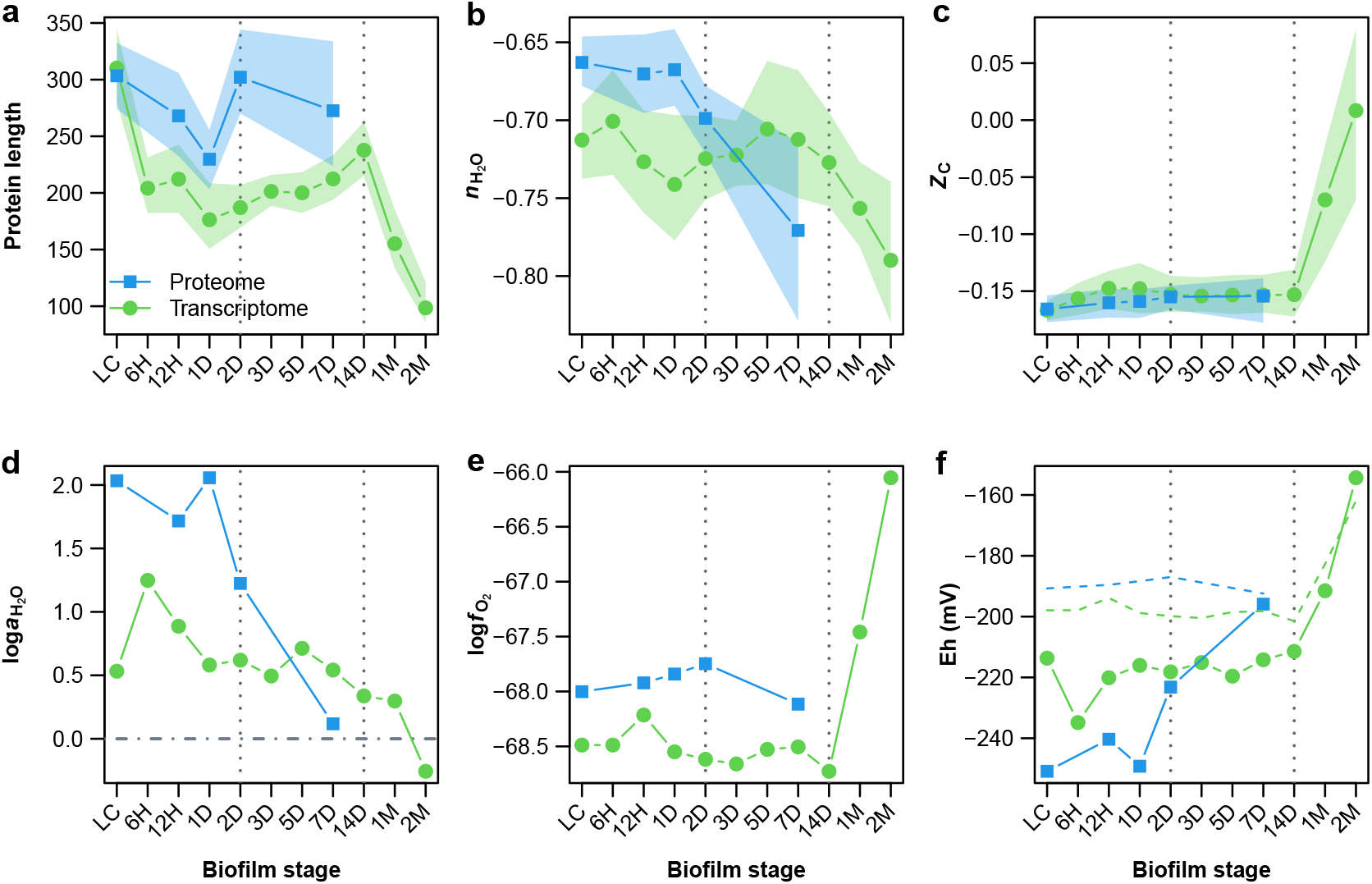
Chemical and thermodynamic analysis of proteins in developing *B. subtilis* biofilms. Dotted lines represent transitions between early and mid periods (2D) and mid and late periods (14D); LC stands for the liquid culture used for inoculation of biofilms. Means for chemical metrics weighted by normalized expression levels of proteins or transcripts (Futo et al., 2021): **a** Protein length; **b** stoichiometric hydration state 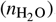; **c** carbon oxidation state (*Z*_C_). The shaded areas represent bootstrap confidence intervals. Thermodynamic parameters computed for target proteins modeled as the mean amino acid compositions for growth stages weighted by abundance of transcripts or proteins: **d** and **e** Optimal values of 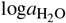 and 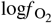, i.e. those that maximize the equilibrium activity of the target proteins against the background proteome for *B. subtilis*. Thermodynamic parameters are mean values for 100 runs, each consisting of 2000 randomly selected background proteins. **f** Virtual redox potential (Eh) computed from Eqs. (3)–(4) and the optimal values of 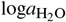 and 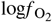. Dashed lines indicate Eh values calculated for the same 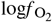values and unit activity of H_2_O

Here, the chemical metrics for individual proteins at each time point were used to compute abundance-weighted mean values of *Z*_C_ and 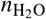. After remaining nearly constant in the early period, the mean 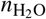of the proteome-based sequences declines in the first transition and mid developmental period (2 to 7 days) (Fig. 5b). The time course for the transcriptome-based abundances exhibits fluctuating 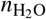of proteins during this time, followed by a steady drop from 7 days to 2 months. Unfortunately, proteomic data for this period are not available for comparison. Through early and mid development there is relatively little variation in mean *Z*_C_ of the proteome, but the transcriptome-based protein sequences become much more oxidized in the late period (Fig. 5c).

To calculate thermodynamic parameters, amino acid compositions of target proteins were calculated as the abundance-weighted means for sequences inferred from transcriptomes and proteomes. The declining trend of 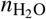is reflected in values of 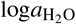 computed from the MaximAct thermodynamic model, which are positive at all but the last time point (Fig. 5d). This can be interpreted as a signal of relatively high hydration potential during the early growth period, followed by a proteomic state that is closer to equilibrium with an aqueous growth medium. The ambient 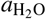 is likely to be not far from unity, as it has been shown that the activity of H_2_O in agar medium is very close to that of the solution used to prepare the medium (Gervais et al., 1988).

Optimal values of 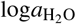 were combined with those of 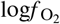(Fig. 5e) to compute virtual Eh using Eqs. (3)–(4). The virtual redox potential rises through development, especially in the last period (Fig. 5f). Even if water activity is set to unity instead of taken from the MaximAct model, the sharp rise in *Z*_C_ for the transcriptome-inferred proteins is associated with a predicted increase in virtual Eh. Notably, the formation of wrinkles in biofilms is postulated to enhance oxygen uptake (Okegbe et al., 2014), and wrinkling in *E. coli* biofilms has been experimentally associated with greater oxygen penetration than into the flat base of the biofilm (Hartmann et al., 2021). The carbon oxidation state of transcriptome-inferred proteins rises in 1-to 2-month-old *B. subtilis* biofilms, well after the appearance of wrinkles in day 3 (Futo et al., 2021). This suggests a delay between wrinkling and the transcriptional adjustment to hypothesized oxidizing conditions within the biofilm. Proteomic and oxygen microelectrode or other redox measurements should be obtained at all growth stages in the same biofilms to enable a direct comparison between this model and experiments.

### Hydration dynamics in development of fruit flies

The fruit fly, *Drosophila melanogaster*, is a well-studied invertebrate model organism in genetics and developmental biology. The changes of whole-organism water content and other biochemical constituents during the development of *D. melanogaster* from larvae to adults, when grown on chemically defined axenic medium, were reported by Church and Robertson (1966). As larvae progress through different instars (i.e. a few days post-hatching), the water content first rises to *>* 80%, then varies between about 75 to 80%. The water content decreases suddenly to 66% in the prepupal stage followed by a small rise in adults (Fig. 6a).

**Fig. 6.**
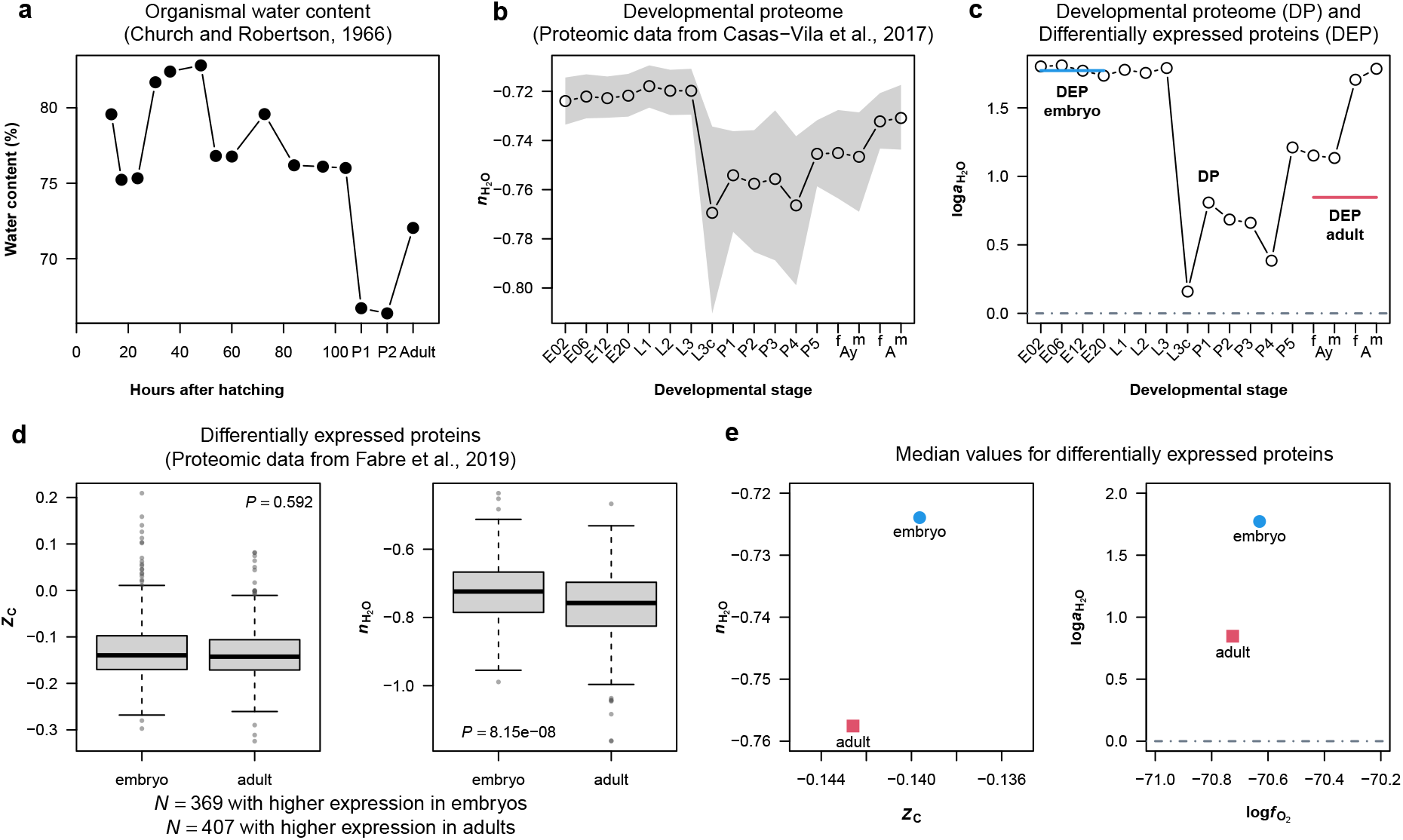
**a** Organismal water content during growth of fruit flies (*D. melanogaster*) from larvae (hours after hatching) to pupal and adult stages, replotted from Figure 4 of Church and Robertson (1966). **b** Mean stoichiometric hydration state of proteins computed from the developmental proteome of Casas-Vila et al. (2017). The shaded area represents the bootstrap confidence intervals. Labels starting with “E”, “L”, and “P” refer to embryogenesis, larval, and pupal stages. “Ay” and “A” stand for young adults and adults, and “f” and “m” indicate female and male adults. **c** Values of 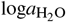 that maximize the activities of target proteins for each developmental time point taken against the *D. melanogaster* proteomic background. Horizontal lines represent values computed for the differentially expressed proteins in embryos and adults using data from a different study (Fabre et al., 2019). **d** Values of *Z*_C_ and 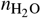computed for differentially expressed proteins in embryo and adult flies (Fabre et al., 2019). The plots were made using R’s boxplot() function; the bottom and top of the boxes correspond to the first and third quartile, the notch (thick bar) indicates the median, and the whiskers are drawn at a data point that is no more than 1.5 times the height of the box away from the box. Individual points beyond the whiskers are plotted. *P*-values were calculated using the two-sided Wilcoxon rank sum test. **e** Median values of *Z*_C_ and 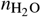from the previous plots and median values of optimal 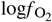and 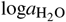 for all differentially expressed proteins in embryos and adults

Casas-Vila et al. (2017) reported proteomic data for developmental stages of *D. melanogaster*, including embryogenesis, larvae, pupae, and adults. The mean stoichiometric hydration state of proteins from the fly developmental proteome is plotted in Fig. 6b. The 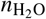is almost constant during embryogenesis and three instars of larvae (L1, L2, L3). Then, 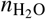drops abruptly in stage L3c (L3 crawling larva). The pupae collected on different days (P1 to P5) exhibit a some-what higher and variable 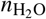, which is still lower than the embryos. The 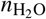increases in young adults and then again in old adults, which have 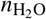values somewhat less than those of embryos and early larvae.

Remarkably, the stoichiometric hydration state of proteins drops precipitously when the crawling larvae leave the medium, at the same time as a sharp drop in organismal water content (Church and Robertson, 1966). Later, a higher organismal water content in adult flies compared to pupae is again reflected in rising proteomic 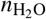for adults. Water activity computed from the MaximAct analysis, in which the target proteins are modeled as having the mean amino acid composition for the proteome at each developmental time point, is plotted in Fig. 6c. The theoretical values of 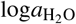 are all positive, with the closest approach to unit activity of H_2_O in stage L3c. The highest values are found for embryos, early larvae, and older adults.

Analysis of proteomic data from a different study (Fabre et al., 2019) shows significantly higher stoichiometric hydration state of proteins enriched in embryos compared to adults, in contrast to indistinguishable mean values of carbon oxidation state (Fig. 6d). For this dataset, only binary classification of differentially expressed proteins was used, and the MaximAct analysis was performed for each differentially expressed protein rather than for mean amino acid compositions. Although the precise age of the adult flies was not given by Fabre et al. (2019), their description refers to the emergence of adults less than 9 days after egg laying. Considering the number of days spent in larval and pupal growth, this suggests an age closer to the young adults (4 hours after eclosure) than the old adults (1 week-old flies) studied by Casas-Vila et al. (2017). Accordingly, the developmental proteome of Casas-Vila et al. (2017) and differentially expressed proteins of Fabre et al. (2019) both exhibit considerably higher modeled water activity in embryos than in young adults (Fig. 6c and e).

## Discussion

### Carbon oxidation state and redox potential

Within a thermodynamic framework, rising *Z*_C_ is theoretically associated with increasing environmental O_2_ concentrations, intracellular redox potential, or both. It is therefore tempting to speculate that the initial rise of *Z*_C_ of proteins that appeared after the origin of the first cells (Fig. 1) might be a geobiochemical record of the oxygenation of Earth’s atmosphere and oceans. The Great Oxidation Event refers to the rise of atmospheric oxygen, originally thought of as a single-step increase between 2.45 and 2.32 billion years ago to 1-40% of present levels (Kump, 2008), but now recognized as an approach to near-modern levels followed by a plunge (Lyons et al., 2014). Therefore, the organic processes required for the origin of cellular organisms at ca. 4290 Mya and for that of eukaryotes at 2101 Mya (divergence times from Kumar et al., 2017) must have taken place under vastly different oxygen regimes.

If Earth’s oxygenation is thermodynamically linked to the oxidation state of proteins, then a rise in *Z*_C_ should be apparent for multiple lineages, not only the human one represented in Fig. 1. To investigate the *Z*_C_ trends in other eukaryotic lineages, additional phylostratigraphic data were analyzed for 31 model organisms representing opisthokonts (fungi, animals, and some protists) studied by Liebeskind et al. (2016b) and for homology groups from 435 fully sequenced animal, plant and fungal species reported by James et al. (2021). For every opisthokont, mean *Z*_C_ of proteins in phylostratigraphic age groups shows an overall increase between cellular organisms (GA 1) and Eukaryota (GA 4) and continues to increase to Opisthokonta (GA 5). For later age groups, the lineages of included organisms diverge, as do their *Z*_C_ trends (Fig. 7a). For homology groups in 435 species, *Z*_C_ of non-transmembrane protein domains increases with a fitted slope of 0.015 to 0.018 Gya_-1_ in plants and animals (Fig. 7b). This is comparable to a rise of ca. 0.019 *Z*_C_ / Gya in the human lineage leading up to the opisthokontal emergence at 1105 Mya (Fig. 7a). Therefore, the carbon oxidation state of multiple eukaryotic lineages rose over an interval spanning the Great Oxidation Event.

**Fig. 7.**
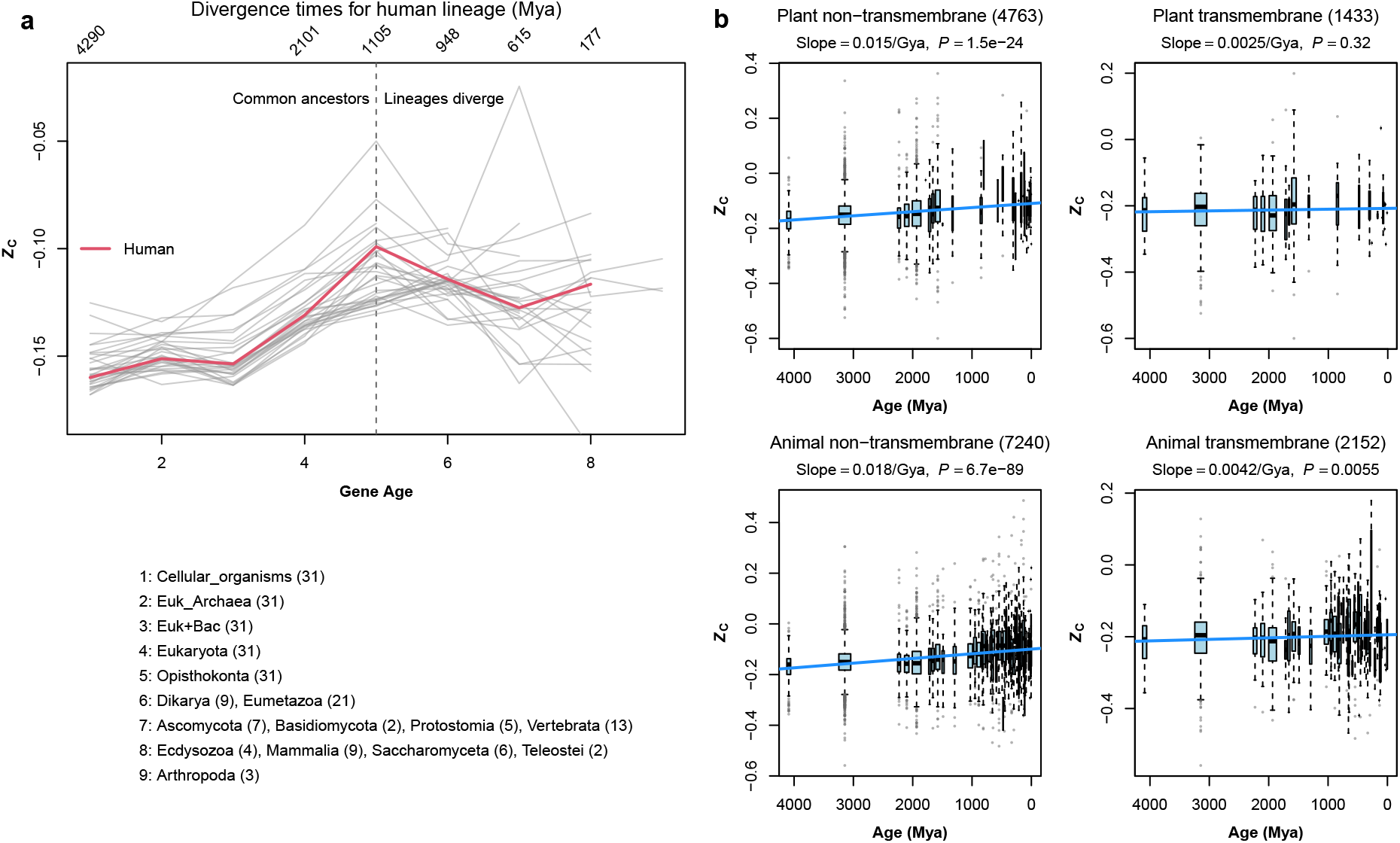
Carbon oxidation state of proteins increases for younger proteins in eukaryotic lineages. **a** Mean protein *Z*_C_ was computed for consensus gene ages of the 31 model organisms in Table 1 of Liebeskind et al. (2016b). All these organisms are opisthokonts (commonly characterized by flagellate cells and formerly known as the Fungi/Metazoa group), so they have a common ancestry up to GA 5, but their lineages diverge for later gene ages. Names of age categories and number of organisms are shown below the plot; the upper labels give the divergence times for the human lineage (Kumar et al., 2017), which is highlighted in red in the plot. **b** *Z*_C_ calculated for amino acid compositions of protein homology groups generated by James et al. (2021). In that study, protein sequences were obtained from 435 genomes, protein domains were annotated using the Pfam database (El-Gebali et al., 2019) and were used to construct homology groups representing the average for each Pfam, and age assignments of Pfam domains were made using TimeTree (Hedges et al., 2006) with exceptions for LUCA and domains that emerged after LUCA but before eukaryotes. Numbers of homology groups are given in the plot titles, and box widths are proportional to the square root of number of groups for each age. Data were obtained from the AAcomp pfam *.csv files in HomologyDictionaryFiles.zip downloaded from Figshare (James et al., 2020)

The last universal common ancestor (LUCA) was most likely an anaerobic chemoautotroph that inhabited reducing hydrothermal environments (Weiss et al., 2016), so it comes as little surprise that the oldest gene ages in Fig. 7a and b correspond to proteins with relatively low carbon oxidation state. The next oldest age group in Fig. 7b is for Pfam domains that emerged after LUCA but before the emergence of eukaryotes from Bacteria and Archaea (as described by James et al., 2021; see also Figure 1 in Weiss et al., 2016). The boxplots in Fig. 7b are consistent with rising *Z*_C_ between protein domains conserved in LUCA and the pre-eukaryotic ancestors. Additional work is needed to confirm whether these trends inferred from amino acid compositions of proteins in extant eukaryotes are also present in the archaeal and bacterial domains.

Compared to transmembrane domains that likely evolved in a hydrophobic environment, non-transmembrane (cytosolic) domains have a higher *Z*_C_ and exhibit a larger and more significant increase of *Z*_C_ across phylostrata (Fig. 7b). A low *Z*_C_ of membrane-associated proteins can be expected based on their high content of hydrophobic amino acids (Dick, 2014). The relatively flat trend for transmembrane domains suggests a smaller evolutionary variability in their elemental composition compared to cytosolic proteins. The findings for cytoplasmic proteins in particular are compatible with a chemical link between the oxygenation of Earth’s atmosphere and the carbon oxidation state of proteins. Rather than being exclusionary, this geobiochemical model is complementary to evolutionary interpretations based on biophysical metrics that are also linked to the amino acid compositions of proteins (e.g. hydrophobicity and intrinsic structural disorder; James et al., 2021).

Following from the chemical analysis, a thermodynamic model allows predicting the virtual redox potentials (Eh) that stabilize target proteins for phylostratigraphic age groups; these Eh values can be compared with biochemical measurements. The Euk Archaea group is of particular interest because it represents the last common ancestor of Eukaryota and Archaea. Although proteins coded by genes in the Euk Archaea group are slightly more oxidized than those of cellular organisms (Fig. 1b), their maximum activity occurs at higher 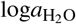. It follows from Eqs. (3)–(4) that the Euk Archaea proteins are stabilized by a lower virtual redox potential, close to −300 mV, which approaches the actual potential of the NADH/NAD_+_ redox couple in mitochondria (Fig. 3f). Notably, such a low virtual redox potential is not possible if the activity of H_2_O is assumed to be unity (dashed line in Fig. 3f). This low virtual redox potential could reflect a reductive overall cellular physiology, typical of archaeal cells, that operated before the endosymbiotic transfer of mitochondria (Martin and Sousa, 2016). The subsequent innovation of oxidative chemistry in aerobes (Williams and Fraústo Da Silva, 2003) is a plausible explanation for the rise in virtual Eh at GA 3 to 5. Further support for this thermodynamic characterization requires analysis of phylostratigraphic data for other lineages, which is not done here.

Although oxygen is required for complex animal life, oxygen concentrations decline rapidly within multicellular structures if the supply is by diffusion alone. The effect of such oxygen gradients on carbon oxidation state of proteins is apparent in laboratory experiments: cells grown in 3D cell culture, whose interiors are characterized by hypoxic conditions, express proteins with lower *Z*_C_ than do cells grown in monolayers (Dick, 2021b). Therefore, a plausible hypothesis is that the beginning of a negative shift in *Z*_C_ around the emergence of Eumetazoa (Figs. 1 and 7a) reflects oxygen gradients in early multicellular structures. The continued drop in subsequent phylostrata might result from hypoxic microniches (for example, those that sustain stem cells) even as circulatory systems became more advanced.

Although speculative at the moment, this type of reasoning centered on oxidation-reduction conditions might one day lead to deeper links between protein evolution and cell physiology.

### Stoichiometric hydration state and water activity

The analysis of stoichiometric hydration state leads to the novel suggestion that protein expression at the proteome scale is chemically coupled to water content in developing organisms. Specifically, for *Drosophila melanogaster*, the large decrease in proteomic 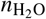at the crawling larva stage (L3c) is aligned with the measured prepupal decrease in organismal water content (Church and Robertson, 1966). Stoichiometric hydration state of proteins also decreases during growth of *Bacillus subtilis* biofilms, but no experimental reports of measured water content could be found for comparison. This chemical representation suggests that the hydration dynamics of developing biofilms might share similarities with that of animals, for which a large body of work documents decreasing water content through early developmental stages (Logan and Himwich, 1972; Calcagno et al., 1972; Friis-Hansen, 1983). Together with the relatively high 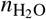of proteins in the earliest phylostratigraphic age groups, these results can be used to postulate that elevated hydration potential is a key factor in the maintenance of protein expression patterns that are characteristic of both embryos and unicellular organisms.

Thermodynamic models in geochemistry often involve one or more “perfectly mobile components”, which are represented by their chemical potentials, instead of by bulk composition (Rumble, 1982; Evans et al., 2013). Oxygen fugacity is one such descriptive variable in many geochemical models, while water activity is often considered to remain close to unity except for certain systems such as brines. In contrast, in this study the chemical potentials of both O_2_ and H_2_O were used as independent variables in order to explore the landscape of relative protein stabilities in two dimensions. The theoretical 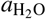 reaches much lower values than measurements in saturated salt solutions (e.g. saturation of NaCl corresponds to 0.755 water activity; Stevenson et al., 2015). At the other extreme, the theoretical values can be greater than unity. This represents an unphysical condition as pure water has unit activity. It may be possible to obtain 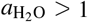 in molecular dynamics simulations of mixtures of H_2_O and organic media due to oversaturation of H_2_O and cluster formation in a nonpolar solvent, but those results were regarded as anomalous by the investigators (Wedberg et al., 2012).

By analogy with the situation for oxygen fugacity in petrology, where vanishingly low values have no physical meaning as partial pressure but remain useful as indicators of oxidation potential (Frost, 1991), the theoretical range of water activity in this study must be interpreted as a theoretical indicator of hydration potential that may not coincide with experimental measurements. Despite this, it is noteworthy that the thermodynamic analysis yields theoretical values of 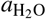 that approach unity for later phylostratigraphic age groups and for advanced stages of biofilm development (Fig. 3e and Fig. 5d). An exception seems to be stages of fly development, which are poised at higher water activity (Fig. 6c). This could be in part a consequence of higher 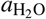 predicted by using the background proteome from *Drosophila* compared to those from other organisms. A productive line of inquiry would be to find a universal set of background proteins with the property that thermodynamic models for various evolutionary and developmental datasets overlap with unit water activity, that is, a physically realistic condition.

In summary, the amino acid compositions and abundances of proteins in principle reflect a water-dominated but highly dynamic hydration state of cells. Besides its importance for polymerization reactions in prebiotic conditions and for macromolecular crowding and phase separation in living cells (Bagatolli and Stock, 2021), water activity can also be regarded as a fundamental thermodynamic parameter for organismal development and Darwinian evolution.

## Declarations

### Availability of data and material

Data were obtained from the UniProt database (The UniProt Consortium, 2019) and the supporting files of previous studies (Liebeskind et al., 2016b; Casas-Vila et al., 2017; Trigos et al., 2017; Fabre et al., 2019; Futo et al., 2021; James et al., 2021) as described in the Materials and Methods and figure captions. Files with phylostrata assignments and UniProt IDs are available in the R package canprot version 1.1.0 (Dick, 2021b). Processed data files used in this study are in the “extdata/evdevH2O” directory of the JMDplots package (version 1.2.10 deposited on Zenodo with accession number 5811079) (Dick, 2021a), except for the differential expression dataset of Fabre et al. (2019), which is located under “extdata/expression” and “extdata/aa”.

### Code availability

The code used to make the figures for this paper is available in the JMDplots package (Dick, 2021a). The maximum activity analysis described here has been implemented in a new function named MaximAct(). The “evdevH2O.Rmd” vignette in the package runs the functions to make each of the figures.

## References

Anderson GM (2005) Thermodynamics of Natural Systems, 2nd edn. Cambridge University Press, Cambridge, doi:10.1017/CBO9780511840258

Anderson GM, Crerar DA (1993) Thermodynamics in Geochemistry: The Equilibrium Model. Oxford University Press, New York, doi:10.1093/oso/9780195064643.001.0001

Bagatolli LA, Stock RP (2021) Lipids, membranes, colloids and cells: A long view. Biochim Biophys Acta, Biomembr 1863(10):183684, doi:10.1016/j.bbamem.2021.183684

Bard JP (1986) Microtextures of Igneous and Metamorphic Rocks. D. Reidel, Dordrecht, Holland, doi:10.1007/978-94-009-4640-8

Brocchieri L, Karlin S (2005) Protein length in eukaryotic and prokaryotic proteomes. Nucleic Acids Res 33(10):3390–3400, doi:10.1093/nar/gki615

Calcagno PL, Hollerman CE, Jose PA (1972) Total body water: Man. In: Altman PL, Dittmer DS (eds) Biology Data Book, vol 3, 2nd edn, Federation of American Societies for Experimental Biology, Bethesda, Maryland, pp 1986–1989

Canty A, Ripley BD (2021) boot: Bootstrap R (S-Plus) Functions. R package version 1.3-28

Capra JA, Stolzer M, Durand D, Pollard KS (2013) How old is my gene? Trends Genet 29(11):659–668, doi:10.1016/j.tig.2013.07.001

Casas-Vila N, Bluhm A, Sayols S, Dinges N, Dejung M, Altenhein T, Kappei D, Altenhein B, Roignant JY, Butter F (2017) The developmental proteome of Drosophila melanogaster. Genome Res 27(7):1273–1285, doi:10.1101/gr.213694.116

Catling DC, Claire MW (2005) How Earth’s atmosphere evolved to an oxic state: A status report. Earth Planet Sci Lett 237(1):1–20, doi:10.1016/j.epsl.2005.06.013

Church RB, Robertson FW (1966) A biochemical study of the growth of Drosophila melanogaster. J Exp Zool 162(3):337–351, doi:10.1002/jez.1401620309

Cramer W (1916) On the biochemical mechanism of growth. J Physiol 50(5):322–334, doi:10.1113/jphysiol.1916.sp001758

Davison AC, Hinkley DV (1997) Bootstrap Methods and Their Application. Cambridge University Press, Cambridge, doi:10.1017/CBO9780511802843

Denbigh K (1981) The Principles of Chemical Equilibrium, 4th edn. Cambridge University Press, Cambridge, doi:10.1017/CBO9781139167604

Dick JM (2014) Average oxidation state of carbon in proteins. J R Soc Interface 11:20131095, doi:10.1098/rsif.2013.1095

Dick JM (2019) CHNOSZ: Thermodynamic calculations and diagrams for geochemistry. Front Earth Sci 7:180, doi:10.3389/feart.2019.00180

Dick JM (2021a) JMDplots 1.2.10. Zenodo, doi:10.5281/zenodo.5811079

Dick JM (2021b) Water as a reactant in the differential expression of proteins in cancer. Comput Syst Oncol 1(1):e1007, doi:10.1002/cso2.1007

Dick JM, LaRowe DE, Helgeson HC (2006) Temperature, pressure, and electrochemical constraints on protein speciation: Group additivity calculation of the standard molal thermodynamic properties of ionized unfolded proteins. Biogeosciences 3(3):311–336, doi:10.5194/bg-3-311-2006

Dick JM, Yu M, Tan J (2020) Uncovering chemical signatures of salinity gradients through compositional analysis of protein sequences. Biogeosciences 17(23):6145–6162, doi:10.5194/bg-17-6145-2020

Domazet-Lošo T, Carvunis AR, Albà MM, Šestak MS, Bakarić R, Neme R, Tautz D (2017) No evidence for phylostratigraphic bias impacting inferences on patterns of gene emergence and evolution. Mol Biol Evol 34(4):843–856, doi:10.1093/molbev/msw284

Downing JE, Christopherson WM, Broghamer WL (1962) Nuclear water content during carcinogenesis. Cancer 15(6):1176–1180, doi:10.1002/1097-0142(196211/12)15:6<1176::AID-CNCR2820150614>3.0.CO;2-F

El-Gebali S, Mistry J, Bateman A, Eddy SR, Luciani A, Potter SC, Qureshi M, Richardson LJ, Salazar GA, Smart A, Sonnhammer ELL, Hirsh L, Paladin L, Piovesan D, Tosatto SCE, Finn RD (2019) The Pfam protein families database in 2019. Nucleic Acids Res 47(D1):D427–D432, doi:10.1093/nar/gky995

Evans KA, Powell R, Frost BR (2013) Using equilibrium thermodynamics in the study of metasomatic alteration, with an application to serpentinites. Lithos 168-169:67–84, doi:10.1016/j.lithos.2013.01.016

Fabre B, Korona D, Lees JG, Lazar I, Livneh I, Brunet M, Orengo CA, Russell S, Lilley KS (2019) Comparison of Drosophila melanogaster embryo and adult proteome by SWATH-MS reveals differential regulation of protein synthesis, degradation machinery, and metabolism modules. J Proteome Res 18(6):2525–2534, doi:10.1021/acs.jproteome.9b00076

Foley SF (2011) A reappraisal of redox melting in the Earth’s mantle as a function of tectonic setting and time. J Petrol 52(7-8):1363–1391, doi:10.1093/petrology/egq061

Frenkel-Pinter M, Rajaei V, Glass JB, Hud NV, Williams LD (2021) Water and life: The medium is the message. J Mol Evol 89(1):2–11, doi:10.1007/s00239-020-09978-6

Friis-Hansen B (1983) Water distribution in the foetus and newborn infant. Acta Paediatr 72(305):7–11, doi:10.1111/j.1651-2227.1983.tb09852.x

Frost BR (1991) Introduction to oxygen fugacity and its petrologic importance. In: Oxide Minerals, Reviews in Mineralogy, vol 25, De Gruyter, pp 1–10, doi:10.1515/9781501508684-004

Futo M, Opašić L, Koska S, Čorak N, Široki T, Ravikumar V, Thorsell A, Lenuzzi M, Kifer D, Domazet-Lošo M, Vlahoviček K, Mijakovic I, Domazet-Lošo T (2021) Embryo-like features in developing Bacillus subtilis biofilms. Mol Biol Evol 38(1):31–47, doi:10.1093/molbev/msaa217

Garrels RM, Christ CL (1965) Solutions, Minerals, and Equilibria. Harper & Row, New York

Gervais P, Molin P, Grajek W, Bensoussan M (1988) Influence of the water activity of a solid substrate on the growth rate and sporogenesis of filamentous fungi. Biotechnol Bioeng 31(5):457–463, doi:10.1002/bit.260310510

Haar L, Gallagher JS, Kell GS (1984) NBS/NRC Steam Tables: Thermodynamic and Transport Properties and Computer Programs for Vapor and Liquid States of Water in SI Units. Hemisphere Publishing Corporation, Washington, D. C.

Hartmann R, Jeckel H, Jelli E, Singh PK, Vaidya S, Bayer M, Rode DKH, Vidakovic L, Díaz-Pascual F, Fong JCN, Dragoš A, Lamprecht O, Thöming JG, Netter N, Häussler S, Nadell CD, Sourjik V, Kovács ÁT, Yildiz FH, Drescher K (2021) Quantitative image analysis of microbial communities with BiofilmQ. Nat Microbiol 6(2):151–156, doi:10.1038/s41564-020-00817-4

Heames B, Schmitz J, Bornberg-Bauer E (2020) A continuum of evolving de novo genes drives protein-coding novelty in Drosophila. J Mol Evol 88(4):382–398, doi:10.1007/s00239-020-09939-z

Hedges SB, Dudley J, Kumar S (2006) TimeTree: A public knowledge-base of divergence times among organisms. Bioinformatics 22(23):2971–2972, doi:10.1093/bioinformatics/btl505

Huang H, McGarvey PB, Suzek BE, Mazumder R, Zhang J, Chen Y, Wu CH (2011) A comprehensive protein-centric ID mapping service for molecular data integration. Bioinformatics 27(8):1190–1191, doi:10.1093/bioinformatics/btr101

James J, Willis S, Nelson P, Weibel C, Kosinski L, Masel J (2020) Data from: Universal and taxon-specific trends in protein sequences as a function of age. Figshare, doi:10.6084/m9.figshare.12037281.v1

James JE, Willis SM, Nelson PG, Weibel C, Kosinski LJ, Masel J (2021) Universal and taxon-specific trends in protein sequences as a function of age. eLife 10:e57347, doi:10.7554/eLife.57347

Jones DP, Sies H (2015) The redox code. Antioxid Redox Signaling 23(9):734–746, doi:10.1089/ars.2015.6247

Kasting JF (1993) Earth’s early atmosphere. Science 259(5097):920–926, doi:10.1126/science.11536547

Kinniburgh DG, Cooper DM (2004) Predominance and mineral stability diagrams revisited. Environ Sci Technol 38(13):3641–3648, doi:10.1021/es034927l

Kitadai N (2014) Thermodynamic prediction of glycine polymerization as a function of temperature and pH consistent with experimentally obtained results. J Mol Evol 78(3-4):171–187, doi:10.1007/s00239-014-9616-1

Kumar S, Stecher G, Suleski M, Hedges SB (2017) TimeTree: A resource for timelines, timetrees, and divergence times. Mol Biol Evol 34(7):1812–1819, doi:10.1093/molbev/msx116

Kump LR (2008) The rise of atmospheric oxygen. Nature 451(7176):277–278, doi:10.1038/nature06587

Lamadrid HM, Rimstidt JD, Schwarzenbach EM, Klein F, Ulrich S, Dolocan A, Bodnar RJ (2017) Effect of water activity on rates of serpentinization of olivine. Nat Commun 8(1):16107, doi:10.1038/ncomms16107

LaRowe DE, Dick JM (2012) Calculation of the standard molal thermodynamic properties of crystalline peptides. Geochim Cosmochim Acta 80:70–91, doi:10.1016/j.gca.2011.11.041

Liebeskind B, McWhite CD, Hines K (2016a) Gene-Ages v1.0. Zenodo, doi:10.5281/zenodo.51708

Liebeskind BJ, McWhite CD, Marcotte EM (2016b) Towards consensus gene ages. Genome Biol Evol 8(6):1812–1823, doi:10.1093/gbe/evw113

Lipman DJ, Souvorov A, Koonin EV, Panchenko AR, Tatusova TA (2002) The relationship of protein conservation and sequence length. BMC Evol Biol 2(1):20, doi:10.1186/1471-2148-2-20

Logan JE, Himwich WA (1972) Animal tissues and organs: Water content. In: Altman PL, Dittmer DS (eds) Biology Data Book, vol 1, 2nd edn, Federation of American Societies for Experimental Biology, Bethesda, Maryland, pp 392–398

Lyons TW, Reinhard CT, Planavsky NJ (2014) The rise of oxygen in Earth’s early ocean and atmosphere. Nature 506(7488):307–315, doi:10.1038/nature13068

Martin WF, Sousa FL (2016) Early microbial evolution: The age of anaerobes. Cold Spring Harbor Perspect Biol 8(2), doi:10.1101/cshperspect.a018127

McIntyre GI (2006) Cell hydration as the primary factor in carcinogenesis: A unifying concept. Med Hypotheses 66(3):518–526, doi:10.1016/j.mehy.2005.09.022

Moyers BA, Zhang J (2017) Further simulations and analyses demonstrate open problems of phylostratigraphy. Genome Biol Evol 9(6):1519–1527, doi:10.1093/gbe/evx109

do Nascimento Vieira A, Kleinermanns K, Martin WF, Preiner M (2020) The ambivalent role of water at the origins of life. FEBS Lett 594(17):2717–2733, doi:10.1002/1873-3468.13815

Natsidis P, Kapli P, Schiffer PH, Telford MJ (2021) Systematic errors in orthology inference and their effects on evolutionary analyses. iScience 24(2):102110, doi:10.1016/j.isci.2021.102110

Okegbe C, Price-Whelan A, Dietrich LEP (2014) Redox-driven regulation of microbial community morphogenesis. Curr Opin Microbiol 18:39–45, doi:10.1016/j.mib.2014.01.006

Olmstead EG (1966) Mammalian Cell Water. Lea & Febiger, Philadelphia

Pace NR (1991) Origin of life: Facing up to the physical setting. Cell 65(4):531–533, doi:10.1016/0092-8674(91)90082-A

R Core Team (2021) R: A Language and Environment for Statistical Computing. R Foundation for Statistical Computing, Vienna, Austria, URL https://www.R-project.org

Ross KFA, Gordon RE (1982) Water in malignant tissue, measured by cell refractometry and nuclear magnetic resonance. J Microsc 128(1):7–21, doi:10.1111/j.1365-2818.1982.tb00433.x

Rumble D III (1982) The role of perfectly mobile components in metamorphism. Annu Rev Earth Planet Sci 10(1):221–233, doi:10.1146/annurev.ea.10.050182.001253

Saryan LA, Hollis DP, Economou JS, Eggleston JC (1974) Nuclear magnetic resonance studies of cancer. IV. Correlation of water content with tissue relaxation times. J Natl Cancer Inst 52(2):599–602, doi:10.1093/jnci/52.2.599

Smith E, Morowitz HJ (2016) The Origin and Nature of Life on Earth. Cambridge University Press, doi:10.1017/CBO9781316348772

Solel E, Tarannam N, Kozuch S (2019) Catalysis: Energy is the measure of all things. Chem Commun 55:5306–5322, doi:10.1039/C9CC00754G

Stevenson A, Cray JA, Williams JP, Santos R, Sahay R, Neuenkirchen N, McClure CD, Grant IR, Houghton JD, Quinn JP, Timson DJ, Patil SV, Singhal RS, Anton J, Dijksterhuis J, Hocking AD, Lievens B, Rangel DEN, Voytek MA, Gunde-Cimerman N, Oren A, Timmis KN, McGenity TJ, Hallsworth JE (2015) Is there a common water-activity limit for the three domains of life? ISME J 9(6):1333–1351, doi:10.1038/ismej.2014.219

Tcherkas YV, Denisenko AD (2001) Simultaneous determination of several amino acids, including homocysteine, cysteine and glutamic acid, in human plasma by isocratic reversed-phase high-performance liquid chromatography with fluorimetric detection. J Chromatogr A 913(1-2):309–313, doi:10.1016/S0021-9673(00)01201-2

The UniProt Consortium (2019) UniProt: A worldwide hub of protein knowledge. Nucleic Acids Res 47(D1):D506–D515, doi:10.1093/nar/gky1049

Trigos AS, Pearson RB, Papenfuss AT, Goode DL (2017) Altered interactions between unicellular and multicellular genes drive hallmarks of transformation in a diverse range of solid tumors. Proc Natl Acad Sci 114(24):6406–6411, doi:10.1073/pnas.1617743114

Van Oss SB, Carvunis AR (2019) De novo gene birth. PLOS Genet 15(5):e1008160, doi:10.1371/journal.pgen.1008160

van ‘t Erve TJ, Wagner BA, Ryckman KK, Raife TJ, Buettner GR (2013) The concentration of glutathione in human erythrocytes is a heritable trait. Free Radic Biol Med 65:742–749, doi:10.1016/j.freeradbiomed.2013.08.002

Wagman DD, Evans WH, Parker VB, Schumm RH, Halow I, Bailey SM, Churney KL, Nuttall RL (1982) The NBS tables of chemical thermodynamic properties. Selected values for inorganic and C1 and C2 organic substances in SI units. J Phys Chem Ref Data 11(Suppl. 2):1–392, URL https://srd.nist.gov/JPCRD/jpcrdS2Vol11.pdf

Wedberg R, Abildskov J, Peters GH (2012) Protein dynamics in organic media at varying water activity studied by molecular dynamics simulation. J Phys Chem B 116(8):2575–2585, doi:10.1021/jp211054u

Weiss MC, Sousa FL, Mrnjavac N, Neukirchen S, Roettger M, Nelson-Sathi S, Martin WF (2016) The physiology and habitat of the last universal common ancestor. Nat Microbiol 1:16116, doi:10.1038/nmicrobiol.2016.116

Williams RJP, Fraústo Da Silva JJR (2003) Evolution was chemically constrained. J Theoret Biol 220(3):323–343, doi:10.1006/jtbi.2003.3152

Wilson BA, Foy SG, Neme R, Masel J (2017) Young genes are highly disordered as predicted by the preadaptation hypothesis of de novo gene birth. Nat Ecol Evol 1(6):0146, doi:10.1038/s41559-017-0146

Winzler RJ (1959) The chemistry of cancer tissue. In: Homburger F (ed) The Physiopathology of Cancer, 2nd edn, Hoeber-Harper, New York, pp 686–706

Zhou JX, Cisneros L, Knijnenburg T, Trachana K, Davies P, Huang S (2018) Phylostratigraphic analysis of tumor and developmental transcriptomes reveals relationship between oncogenesis, phylogenesis and ontogenesis. Converg Sci Phys Oncol 4(2):025002, doi:10.1088/2057-1739/aab1b0

